# Making the most out of it: shallow genome-skimming possibilities for the systematics of prickly lineages of *Solanum* (Solanaceae)

**DOI:** 10.64898/2026.07.08.737304

**Authors:** Richard Tarcisio de Lima Alves, Yuri Fernandes Gouvêa, Jeronymo Dalapicolla, Peter Poczai, Leandro Lacerda Giacomin

## Abstract

**Premise:** Genome skimming (GS) is a cost-effective approach for plant phylogenomics, but its ability to recover informative datasets from different genomic compartments, particularly genome-wide SNPs, remains poorly explored in *Solanum*.

**Methods:** We evaluated shallow GS for phylogenetic inference in South American prickly *Solanum* lineages by recovering plastid, mitochondrial, and nuclear datasets, including coding regions and genome-wide SNPs. Phylogenies were inferred using maximum-likelihood and coalescent approaches under different SNP filtering strategies.

**Results:** GS successfully recovered complete plastomes, organellar coding regions, and large SNP datasets, but failed to consistently assemble mitochondrial genomes or recover low-copy nuclear genes. SNP-based analyses, especially from the nuclear genome, produced stable, well-supported phylogenies that were largely congruent across inference methods. In contrast, coding-region datasets, particularly from the mitochondrial genome, showed greater topological discordance, revealing cytonuclear conflict.

**Discussion:** Our results demonstrate that shallow GS is an effective strategy for generating informative SNP datasets for phylogenetic inference in *Solanum*, despite limitations in recovering complete mitochondrial genomes and low-copy nuclear loci. SNP-based analyses substantially expand the phylogenetic potential of GS, providing a practical and cost-effective alternative for systematic studies.

---

With the advent of high-throughput sequencing (or Next-generation sequencing; NGS) and its significant impact on evolutionary studies in plants, several genome partitioning and capture strategies have emerged as powerful tools in plant genomics (Yu et al., 2018). Among NGS-based approaches, genome skimming (GS) is one of the simplest methodologies, involving random sampling of a small percentage of total genomic DNA (gDNA) (Dodsworth, 2012). Shallow GS (i.e., up to 5x the genome size) generally yields shallow sequencing of the nuclear genome, and relatively deep sequencing of the plastome, mitogenome, and repetitive nuclear sequences (Straub et al., 2012). Genome skimming is a cost and time-effective approach that enables recovery of organellar genomes and repetitive nuclear regions, supporting phylogenomic inference even under low coverage (Straub et al., 2011; Bakker et al., 2016; Chen et al., 2024).

Most plant phylogenomic studies have focused on plastid data (e.g., Zhong et al., 2010; Gitzendanner et al., 2018; Li et al., 2021), whereas nuclear and mitochondrial genomes remain comparatively underexplored (e.g., Burleigh et al., 2011; Zhang and Hong-ma, 2024; Zuntini et al., 2024). In particular, SNP-based phylogenomic inference from genome skimming data is still uncommon in plants, despite recent studies demonstrating its potential for recovering robust phylogenetic relationships (e.g., Liu et al., 2014; Lin et al., 2022).

*Solanum* is the largest genus of Solanaceae, comprising approximately 1,250 species, many of them concentrated in the Neotropics (Hilgenhof et al., 2023). Despite major phylogenomic advances, relationships among several lineages of the Leptostemonum clade, the most speciose, remain poorly investigated. Increasing genomic sampling may help resolve these relationships, but large-scale phylogenomic approaches are often costly. Genome skimming combined with SNP recovery may provide a practical alternative for addressing these issues. Important studies have advanced *Solanum* phylogenomics using plastome and nuclear genome data (e.g., Gagnon et al., 2022, Messeder et al., 2024), while the mitogenome has been minimally or virtually not explored for phylogenetic inference in *Solanum*. Furthermore, only one study using SNPs is available for the genus, aiming to reconstruct the phylogeny of *Solanum* section *Petota* Dumort. (Li et al., 2018).

Here, we explore shallow genome skimming approach possibilities for phylogenomics, drawing special attention to the recovery of SNPs to infer phylogenies. For that, we use selected spiny lineages of *Solanum* (Leptostemonum clade of *Solanum*) from South America as a model group, using nuclear, mitochondrial, and plastid genomes across a set of species (Figure 1). Our main goal was to evaluate how genome skimming combined with genetic markers performs in recovering phylogenies within the prickly lineages and the genus as a whole, aiming to upscale approaches for other lineages and plant groups.

**Figure 1:**
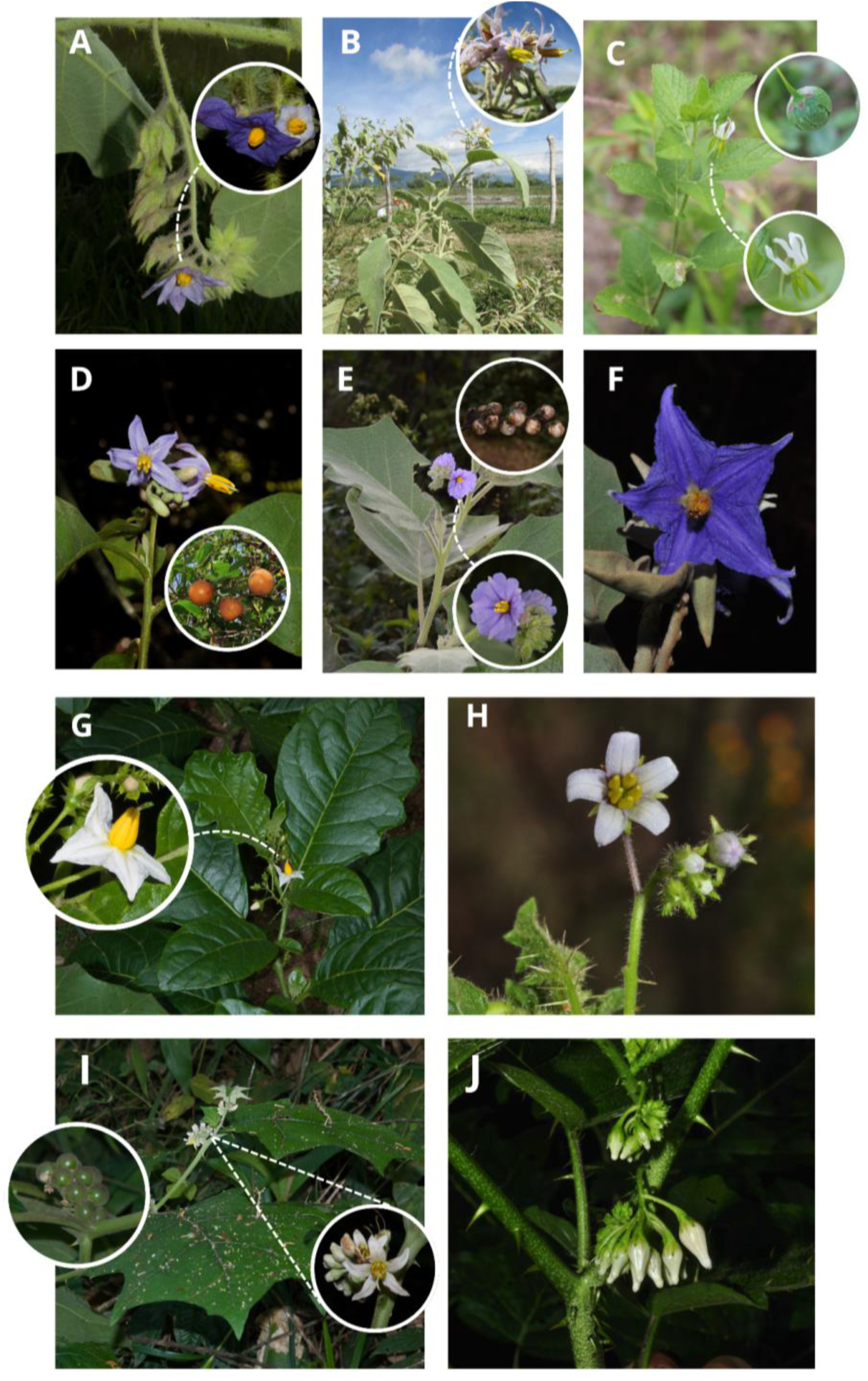
South American spiny species of *Solanum* used in the phylogenetic reconstructions. A: *S. aciculare* Sw., highlighting its lilac-corolla flower and branches densely covered with trichomes (Photo: Y. Gouvea). B: *S. crotonifolium* Dunal., with emphasis on its inflorescence and lilac flowers (Photo: L. Giacomin). C: *S. fernandesii* V. Samp. & R. L. Moura, with emphasis on its white, star-shaped corolla and fruit (Photo: V. Sampaio). D: *S. jussiaei* Dunal., with flowers and fruits highlighted (Photo: Y. Gouvea). E: *S. leptostachys* Sendth., with emphasis on its flower and fruits (Photo: J. Stehmann). F: Lilac flower of *S. lycocarpum* A. St.-Hil. (Photo: Y. Gouvea). G: *S. sessilantherum* Gouvêa & Stehmann, with the white flower highlighted (Photo: Y. Gouvea). H: *S. stenandrum* Sendtn., with emphasis on the white, star-shaped corolla (Photo: L. Giacomin). I: *S. stramoniifolium* Jacq., also highlighting the star-shaped corolla and fruits (Photo: Y. Gouvea). J: *S. vaillantii* Dunal., highlighting the cymose inflorescence and spiny branches (Photo: Y. Gouvea).

## METHODS

### Taxon sampling, DNA extraction and sequencing

To test the different outcomes of genome compartments, we selected the raw GS data for eleven species from the unpublished dataset of Gouvêa (2020) (Table 1). These species represent ten minor neotropical clades within the Leptostemonum clade recognised by Stern et al. (2011) and Gouvêa (2020). These ten neotropical clades are organized into five more comprehensive clades, which we named in this study as: clade A (comprising *Solanum lycocarpum* A.St.-Hil. and *S. leptostachys* Dunal); clade B (*S. crotonifolium* Humb. & Bonpl. ex Dunal and *S. sessilantherum* Gouvêa & Stehmann); clade C (*S. vaillantii* Dunal and *S. stramoniifolium* Jacq.); clade D (*S. stenandrum* Sendtn. and *S. fernandesii* V.Samp. & R.L.Moura); and clade E (*S. jussiaei* Dunal and *S. aciculare* Sw.). We have chosen to use a single closely related outgroup (*S. alternatopinnatum* Steud.), to avoid putative random rooting (DeSalle et al., 2023). Selected species with vouchers, SRA, and number of reads obtained are shown in Table 1.

**Table 1:**
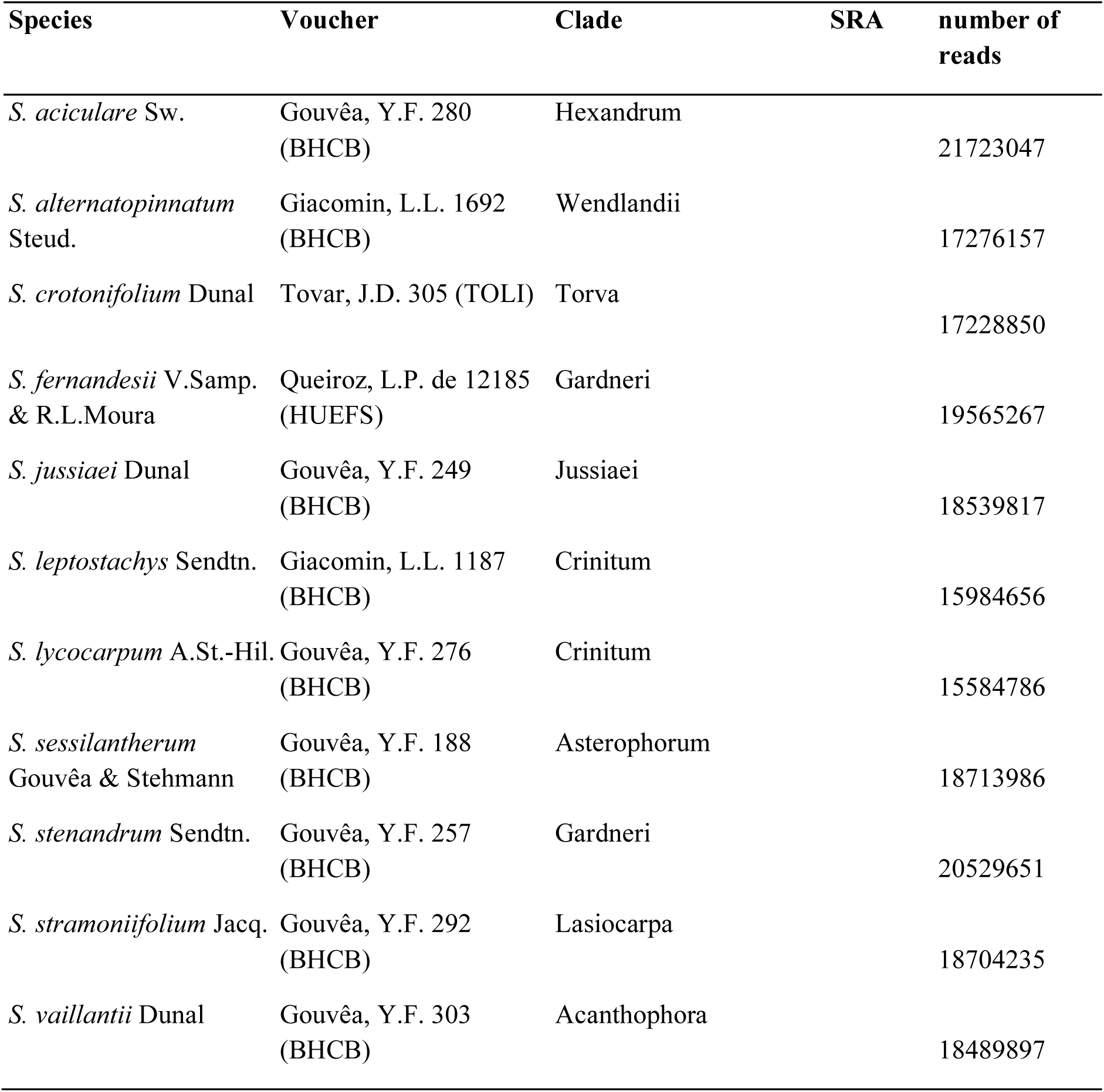
List of *Solanum* clade Leptostemonum taxa included in the analysis, with voucher and herbaria, SRA deposit (to be released upon acceptance), and sequencing yield (number of raw reads) obtained in GS.

Using the DNeasy Plant Mini Kit (Qiagen, Valencia, CA, USA), genomic DNA was extracted from up to 40 mg of silica gel-dried fragments of field-acquired leaf samples. To assemble libraries, at least 800 ng of DNA was aliquoted and mechanically sheared in a Bioruptor ultrasonicator (Diagenode Inc.) to obtain 200 to 500 bp fragments. Libraries were prepared using the Meyer and Kircher (2010) protocol, double-indexed (Kircher et al., 2012) and brought to equimolarity prior to sequencing. Sequencing was performed in two independent lanes of an Illumina HiSeq 4000 platform (2 x 150 paired-end reads) for increased heterogeneity, by Novogene Co. (Sacramento, CA, USA). All sequences are available in the SRA/Genbank, under BioProject code (to be released upon acceptance).

### Data quality control

Raw data was processed by first removing exogenous DNA by mapping to contaminant genomes implemented in FastQ Screen (Wingett and Andrews, 2018). Low complexity reads showing multiple hits to multiple genomes were also removed. At a second filtering step, contaminants associated with herbarium specimens or microorganisms living on plant surfaces were removed by creating a custom bowtie2 (Langmead and Salzberg, 2012) indexed database for FastQ Screen from the genomes listed in Bieker et al. (2020). Low-quality bases and adapters were removed with BBDuk implemented in the BBTools package (Bushnell, 2014) by setting a minimum read quality of 20 and a minimum read length of 60 bp. Read quality was assessed with FastQC (Andrews, 2010).

### Plastome assembly, gene capture, and construction of genome-wide variant matrices

To explore the possible outcomes that can be provided by the GS approach, we tried to assemble different datasets for the three genomic compartments: i) full plastome assembly, using *de novo* and referenced methods; ii) capture of coding plastome regions; iii) assembly of a plastome wide SNP matrix; iv) full mitochondrial genome assembly, using *de novo* and referenced methods; v) capture of coding mitochondrial genome regions; vi) assembly of a mitochondrial genome wide SNP matrix; vii) capture of low-copy universal nuclear gene sets; viii) assembly of a nuclear genome-wide SNP matrix.

Plastomes were assembled using *de novo* assembly with GetOrganelle 1.6.2.e (Jin et al., 2020), and reference-guided assembly using mapping strategy in Geneious R9 (Biomatters Ltd, Auckland, New Zealand), considering the plastome of *S. dulcamara* L. (GenBank KY863443; Amiryousefi et al., 2018) as reference. For the reference-guided approach, regions with error probability higher than 5% were removed prior to assembly, consensus sequences of aligned reads were matched to a minimum coincidence of 50% with the aligned reads, and regions with coverage lower than 3x were trimmed. Contigs of both methods were compared, and no relevant differences were found (mean differences of 0.000013% across species, representing 3 bp). The *de novo* and referenced contigs were superimposed as a consensus, when a full circular structure was not assembled in the *de novo* approach. Assembled plastomes were annotated using the Geneious “transfer annotations” tool by aligning to the previously annotated plastome of *S. dulcamara*, with a similarity threshold of 70%. The same approaches were used to assemble the mitochondrial genome, using the *S. tuberosum* L. mitogenome published by Varré et al. (2019) (GenBank accessions MN114537, MN114538, MN114539), but no structured contig to any of the three recognized molecules could be successfully recovered for all the species. The contigs of the mitochondrial assembly were not used in the downstream analysis, therefore.

To explore the usage of the coding regions of the organelles, a protein-coding gene matrix was assembled for both organelles. Forty mitochondrial and seventy-five chloroplast protein-coding genes were selected from the respective reference genomes and extracted using Geneious Prime. These gene sequences were used as target references for read mapping and assembly via HybPiper v.2.3.2 (Johnson et al., 2016). Target sequence recovery in HybPiper involves three sequential steps: first, raw reads were mapped to the reference gene targets using BWA v.0.7.17 (Li and Durbin, 2009) and sorted with samtools (Danecek et al., 2021). For assembly, we modified the default minimum coverage threshold of SPAdes v.3.15.5 (Bankevich et al., 2012) by lowering the --cov_cutoff to 5, which allowed for the retention of more contigs per gene (Liu et al., 2021). Assembled reads for each gene were then merged into “stitched contigs” using Exonerate v.2.4.0 (Slater and Birney, 2005), aligning them to their corresponding target references.

We also tentatively captured coding regions of the nuclear genome, using specific target files of selected gene sets (Angiosperms353 universal bait set; COSI and COSII genes and ribosomal DNA). The following reference targets were used: i) for the nuclear Angiosperms353 universal bait set, the target provided by Johnson et al. (2019) was used; ii) for the COSI and COSII genes, a target from the tomato genome was used (TGC, 2012); iii) for the ribosomal DNA a target from the tomato genome was used (TGC, 2012). To assess gene recovery efficiency across samples, we employed the post-processing utilities hybpiper stats and hybpiper recovery_heatmap, both implemented in HybPiper. Since none of the nuclear gene sets yielded homogeneous captures, these datasets were not used in downstream analysis (see Supplementary Material for heatmaps of each dataset). Comprehensive recovery statistics for each gene and a visual summary of recovery success across samples are provided; nevertheless, for the organellar genes obtained and shown in Appendix S6 and Appendix S5 (See Supporting Information with this article). All analysis was conducted on the Puhti supercomputer at CSC Espoo, Finland.

To generate high-confidence nuclear single-nucleotide polymorphism (SNP) datasets for phylogenomic inference, we implemented a reference-guided variant calling pipeline following the recent protocol developed by Rick et al. (2024). The procedure was further optimized and modified for organelle genomes and skimming data. The high-quality chromosome-scale genome of cultivated tomato (*S. lycopersicum* L.) (TGC, 2012) was chosen for reference mapping. This genome was shown to be highly syntenic with other important Solanaceae lineages and regarded as the currently available highest quality assembly (Gao et al., 2019; Li et al., 2023). Filtered fastq read files were mapped to the tomato genome using Bowtie2, with the --very-sensitive-local option, which balances alignment sensitivity and specificity, especially suitable for fragmented or lower-depth reads typical of genome skimming. Resulting Sequence Alignment Map (SAM) files were converted to binary alignment map (BAM) files and sorted with samtools (Danecek et al., 2021). To ensure high-confidence read alignments and reduce potential artifacts from multi-mapping, we retained only primary alignments with a minimum mapping quality of 20 using samtools view. Reads flagged as secondary alignments (SAM flag 0x100) were excluded, and only those with a MAPQ score ≥ 20 were retained. This filtering step minimized ambiguity in repetitive genomic regions and enhanced the accuracy of variant detection, ensuring the reliability of downstream phylogenomic analysis. Variant calling was performed with bcftools 1.9. mpileup and call (Li et al., 2009), allowing a probabilistic approach for SNP detection (Li, 2011). To account for differences in genome copy number, we applied distinct read depth thresholds during SNP calling. A maximum read depth of 100 was set for nuclear loci to avoid biases introduced by repetitive regions, paralogy, or excessively high local coverage. We used diploid settings (--ploidy 2) and excluded indels (-V indels) from the resulting variant call format file (VCF; Danecek et al., 2011). To annotate population-level variant statistics, we applied bcftools + fill-tags to enrich VCF files with INFO field tags. Variable sites were further filtered with bcftools, setting the minimum based quality (Phred) score (--min-BQ) to 20, retaining only biallelic variants (view -m2 -M2). Genotype-level confidence was evaluated using the Phred-scaled likelihoods (PL) field, keeping only sites where the difference between the second-best and best genotype likelihoods was >10 (PL[0:1] - PL[0:0] > 10), approximating a GQ > 10 threshold. To minimize the effects of linkage disequilibrium among closely spaced variants, the dataset was subjected to a thinning step using vcftools (Daneck et al., 2011). Only one SNP was retained per 500 bp across the genome (--thin 500), ensuring a more uniform and representative distribution of variants, mitigating the potential bias introduced by tight linkage. In a final filtering step, we removed sites with MAF ≤ 0.07 using vcftools to reduce the influence of rare variants and sequencing noise. The filtered VCF file was converted to phylip format using vcf2phylip v2.0 (Ortiz, 2019), coding heterozygous loci in IUPAC format. We created several phylip files to assess the effect of missing data on phylogenetic inference.

A genome-wide SNP matrix was generated for the organellar genomes using the mitochondrial genome of tomato (Kim and Lee, 2018) and the corresponding chloroplast genome (Daniell et al., 2006) as reference sequences. The analytical pipeline followed the same procedures as described above, with the exception that SNP calling was performed under the assumption of haploidy by specifying --ploidy 1 to account for the uniparental inheritance. For plastid and mitochondrial genomes, which are typically present in significantly higher copy numbers, the maximum depth threshold was increased to 200, allowing retention of informative variation without over-filtering true organellar signals while still minimizing potential sequencing artifacts. No thinning was applied to SNPs from the organelle genomes – a decision based on both biological and practical considerations. First, organelle genomes are relatively small and non-recombining, meaning that linkage disequilibrium (LD) among sites is expected to be inherently high across the entire molecule. Thinning would not mitigate LD in this context, and could instead remove valuable phylogenetic signals in datasets where site density is already limited. Second, organelle genomes often exhibit lower sequence variation compared to nuclear data; therefore, retaining all available SNPs maximizes resolution for inferring organellar phylogenies, especially when analyzing closely related taxa.

### Phylogenomic analyses

A plastid phylogenomic analysis protocol was implemented adjusting for motif-based alignment and adjustment suggested by Roestel et al. (2024). Complete plastid genomes were aligned with MAFFT (Katoh and Standley, 2013) implemented in Geneious Prime (Kearse et al., 2012) using default gap parameter settings. One inverted repeat region (IR) was removed to reduce over representation of genes in the analysis. Ambiguous sites were masked using Geneious Tools creating a stripped alignment. An indel coding approach implemented in bad2matrix (Salinas et al., 2024) was applied to the masked alignment since it was shown that the integration of indels may yield supplementary data enhancing resolution and increasing support (Donath and Stadler, 2018; Suvorov et al., 2020) of problematic nodes in phylogenetic inferences (Neumann et al., 2021). All gaps were removed and the alignment was divided to coding and non-coding intergenic spacer regions, which were manually extracted in Geneious Prime. Alignments containing no informative sites were discarded, and the remaining regions were processed through block filtering using BMGE (Criscuolo and Gribaldo, 2010) with an entropy cutoff 0.5 executed through the graphical interface of Galaxy Pasteur (Mareuil et al., 2017). Alignments were concatenated using bad2matrix manually adjusting the generated partition file for the inclusion of indel coding data. The filtered sub-alignments from plastid DNA and indels were used in a partitioned mixed data analysis (Chernomor et al., 2016) with IQ-TREE v2.3.6 (Minh et al., 2020). The optimal partitioning approach was determined using PartitionFinder (Lanfear et al., 2012), employing a greedy strategy that involved merging partitions until no further improvement in the fitting of the model was observed. Ten runs were executed searching for the best-scoring maximum likelihood tree, with 1000 ultrafast bootstrap approximations (Hoang et al., 2018) and the same number of replicates for SH-like approximate likelihood ratio test (Guindon et al., 2010) to assess branch supports. To reduce the risk of overestimating branch supports, a hill-climbing nearest neighbor interchange (-bnni) search was implemented on the corresponding bootstrap alignment. For the nuclear and mitochondrial SNP matrix IQ-TREE runs followed the same protocol without indel coding and partitioning. We also calculated site concordance factors (sCF, sCF1, sCF2) to assess the percentage of decisive sites supporting a branch tree in the IQ-TREE runs (Minh et al., 2020). In addition, for each dataset, we inferred a species tree under the multispecies coalescent (MSC) model using SVDQuartets (Chifman and Kubatko, 2014) as implemented in PAUP 4.0a* (Kubatko and Knowles, 2023). An exhaustive quartet sampling was performed with the Quartet Fiduccia-Mattheyses (QFM) assembly algorithm (Reaz et al., 2014) and species-membership partitioning. Branch support was assessed with 1,000 bootstrap replicates, and a majority-rule consensus tree was generated, collapsing clades with <50% support into polytomies. Results were visualized using the ggtree package (Yu et al., 2016) in R v4.5 (R Core Team, 2025) with a custom script (to be released upon acceptance).

## RESULTS

### Data Recovery Across Genome types

The attempt to recover diverse datasets from different genomes demonstrated that the shallow genome skimming (GS) approach employed is not suitable for assembling the mitochondrial genome, nor for capturing low-copy nuclear genes for systematic studies (Appendix S4). The attempt to capture nuclear genes using the Angiosperms353, COSI, and COSII bait sets, as well as ribosomal genes, did not yield homogeneous datasets for the species investigated (Appendix S1, S2 and S3).

In contrast, the attempt to recover coding regions from the organellar genomes analyzed, as well as the assembly of plastomes, resulted in comprehensive datasets for investigating the systematics of the group. SNP capture also produced considerably long matrices (Table 2), with unrestricted alignments containing 4,444 plastid genome SNPs, 7,519 mitochondrial genome SNPs, and 138,560 nuclear SNPs. However, filters to reduce missing data and to account for physically linked genes considerably reduced the final matrix size for the mitochondrial and nuclear genomes. The plastid genome SNPs underwent only a small reduction when limiting 10% missing data (4.71%), whereas the mitochondrial genome exhibited a moderate reduction (41.1%). For the nuclear genome, however, the reduction was drastic (94.2%) when applying filters for missing data (30,146 SNPs) and for missing data combined with linked genes (8,048 SNPs) (see Table 2).

**Table 2.**
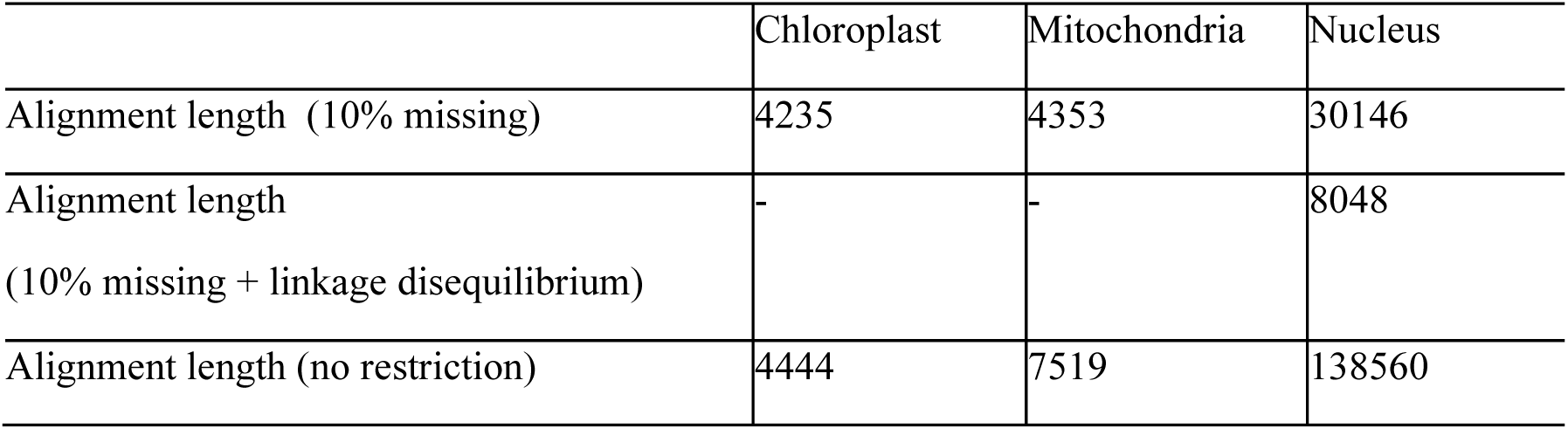
Number of SNPs obtained per genomic compartment with and without missing data restrictions and linkage disequilibrium prunning, limited from 5 to 100x coverage (see Materials and Methods).

|  | Chloroplast | Mitochondria | Nucleus |
| --- | --- | --- | --- |
| Alignment length (10% missing) | 4235 | 4353 | 30146 |
| Alignment length<br>(10% missing + linkage disequilibrium) | - | - | 8048 |
| Alignment length (no restriction) | 4444 | 7519 | 138560 |

### Phylogenetic Hypotheses and Dataset Resolution

The resolution achievable for each dataset obtained is shown through phylogenetic hypotheses of selected neotropical Leptostemonum trees inferred with maximum likelihood (ML) (Figure 2) and coalescent methods (CM) (Figure 3). A summarization of relationships among the clades recovered across the different datasets and methods is presented in Figure 4. The trees inferred with ML showed greater variation in branch and clade support. For example, the mitochondrial tree based on CDS regions (Figure 3F) contained many groupings with low support.

**Figure 2:**
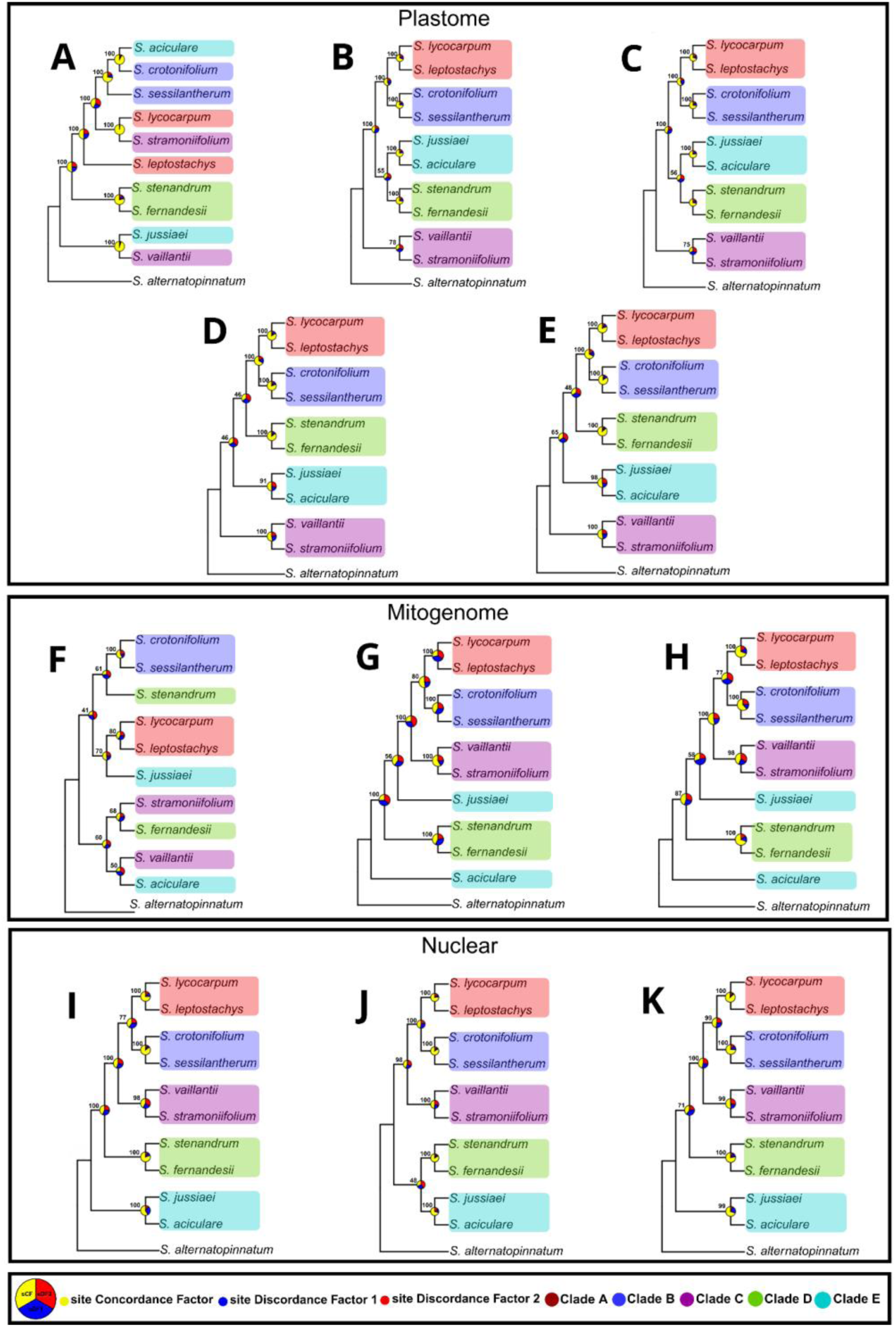
Phylogenetic trees inferred using ML methods. Each tree is derived from a different dataset as follows (see Methods) A. aligned plastid genomes, with the alignment corrected for indels and retaining only one inverted repeat (IR). B. includes plastomes with coding regions (CDS), while panel C displays CDS regions excluding paralogous genes. Panel D presents plastid SNPs including only biallelic loci and no indels, and panel E uses the same dataset but allows up to 10% missing data. Panel F shows the mitochondrial genome with CDS regions. Panel G presents mitochondrial SNPs including only biallelic loci and no indels, and panel H allows up to 10% missing data for the same mitochondrial dataset. Panel I shows nuclear genome SNPs including only biallelic loci and no indels; panel J includes the same nuclear SNP dataset but filtered to allow 10% missing data. Finally, panel K applies a linkage disequilibrium (LD) filter to the nuclear dataset, removing SNPs with high LD.

**Figure 3:**
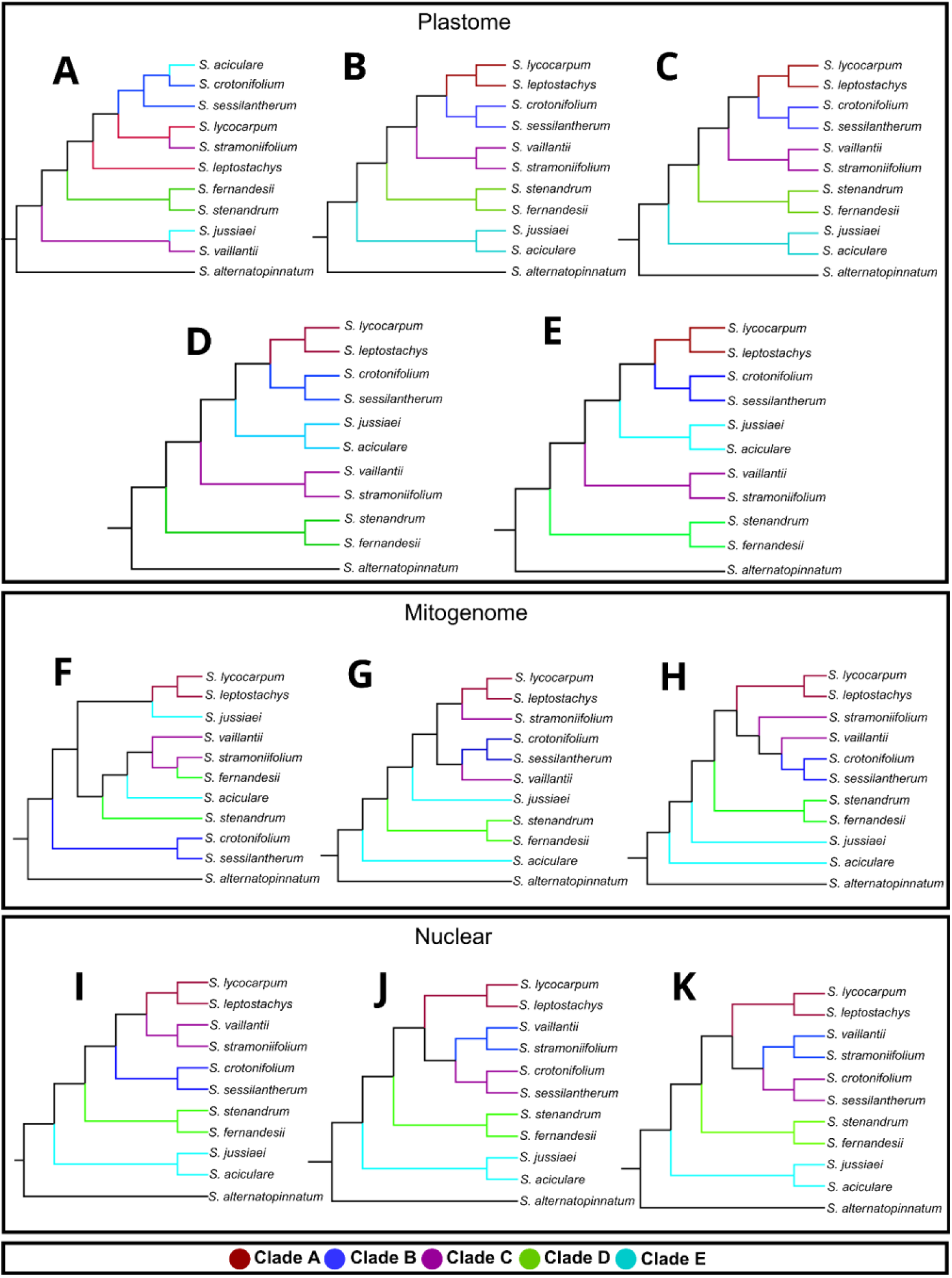
Phylogenetic trees inferred using coalescence methods. All branches have 100% Bootstrap support. Each tree is derived from a different dataset as follows (see Methods) A. aligned plastid genomes, with the alignment corrected for indels and retaining only one inverted repeat (IR). B. includes plastomes with coding regions (CDS), while panel C displays CDS regions excluding paralogous genes. Panel D presents plastid SNPs including only biallelic loci and no indels, and panel E uses the same dataset but allows up to 10% missing data. Panel F shows the mitochondrial genome with CDS regions. Panel G presents mitochondrial SNPs including only biallelic loci and no indels, and panel H allows up to 10% missing data for the same mitochondrial dataset. Panel I shows nuclear genome SNPs including only biallelic loci and no indels; panel J includes the same nuclear SNP dataset but filtered to allow 10% missing data. Finally, panel K applies a linkage disequilibrium (LD) filter to the nuclear dataset, removing SNPs with high LD.

**Figure 4:**
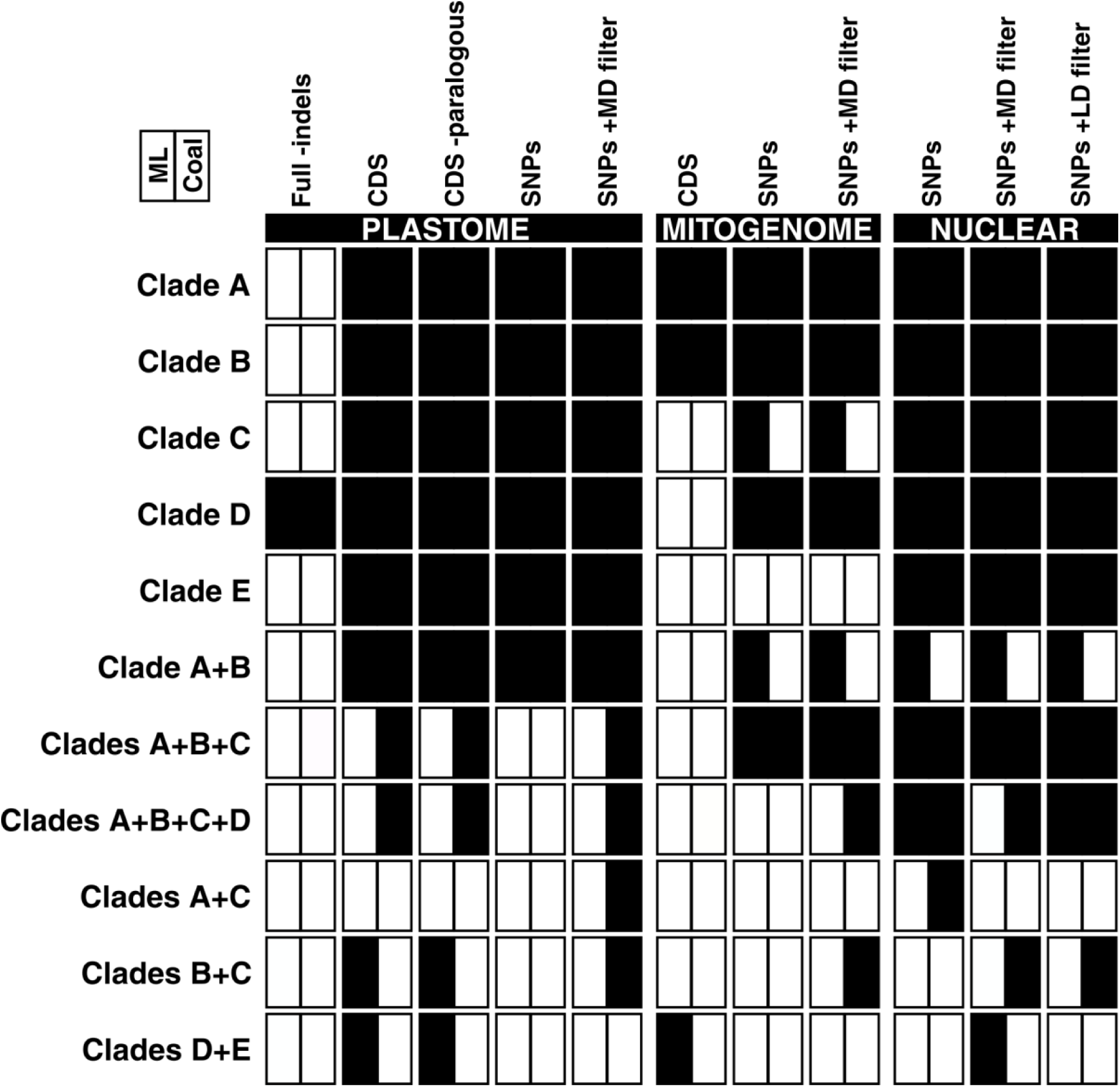
Summary of the main topological differences between phylogenetic trees inferred using maximum likelihood (ML) versus coalescent (Coal) methods across different datasets. Columns represent the datasets used, organized by genome type (see Methods for details), and rows represent *Solanum* clades along with their expected relationships based on previously published studies (see Discussion). Each clade is depicted as a square divided in half: the left half is filled in black if the clade was recovered by ML methods, and the right half only if recovered by coalescent methods. For the trees derived from the plastome, the datasets were: “Full-indels” = aligned plastid genomes, with the alignment corrected for indels and retaining only one inverted repeat (IR). “CDS” = coding regions (CDS) in plastome. “CDS -paralogous” = CDS regions excluding paralogous genes. “SNPs” = plastid SNPs including only biallelic loci and no indels. “SNPs +MD filter” is the same SNPs dataset but allows up to 10% missing data (MD). For the trees derived from the mitogenome, the datasets were: “CDS” = CDS regions from the mitochondrial genome. “SNPs” = mitochondrial SNPs including only biallelic loci and no indels. “ SNPs +MD filter” = the same mitochondrial SNPs dataset but filtered to allow up to 10% missing data (MD). For the nuclear genome: “SNPs” = nuclear genome SNPs including only biallelic loci and no indels. “SNPs +MD filter” = the same nuclear SNP dataset but filtered to allow 10% missing data. “SNPs +LD filter” = the same nuclear SNP dataset but removing SNPs with high linkage disequilibrium (LD), applying a linkage disequilibrium (LD) filter.

Overall, the trees recovered five clades with stable groupings under both inference methods, with some exceptions. The mitochondrial trees, regardless of inference method, displayed the greatest topological differences (see Figures 2F–H and 3).

In topologies recovered using plastome data, the without indels database recovered only one of the five clades (clade D), whereas the other plastome-derived datasets performed similarly regardless of the inference method applied (Figure 4). Topologies inferred with nuclear SNPs recovered all five clades (see below the variations in support, site concordance factors (sCF), and relationships among clades), what was consistent across filtering thresholds used (Figure 2 and Figure 4). For mitogenome-derived datasets, clades A, B, and D were recovered with support. Among these, mitochondrial SNP datasets outperformed those based on mitochondrial coding regions, successfully recovering clade C using ML methods. Overall, mitochondrial SNP-based datasets performed very similarly to nuclear SNP-based ones. In contrast, datasets derived from mitochondrial coding sequences (CDS) showed the second weakest performance in recovering the clades, outperforming only the without Indels in the plastome database (Figure 3).

Clade A comprises *S. lycocarpum* and *S. leptostachys* (corresponding to the Crinitum clade of Stern et al., 2011 and Hilgenhof et al., 2023); Clade B includes *S. crotonifolium* and *S. sessilantherum* (Torva clade of Hilgenhof et al., 2023); Clade C includes *S. vaillantii* and *S. stramoniifolium* (Acanthophora of Stern et al., 2011 and Hilgenhof et al., 2023, and Lasiocarpa of Stern et al., 2011 and Hilgenhof et al., 2023, respectively); Clade D is formed by *S. stenandrum* and *S. fernandesii* (Gardneri clade of Hilgenhof et al., 2023); and finally, Clade E comprises *S. jussiaei* and *S. aciculare* (*S. jussiaei* belonging to Micracantha in Hilgenhof et al., 2023 and *S. aciculare* to Hexandrum in Gouvêa, 2020).

The inferred topologies were similar in most trees, but varied depending on the applied filter. The coalescent plastome tree without indels and with only one inverted repeat (IR) (Figure 3A) differed in some branches and even in species groupings compared to the other topologies, being the most discordant dataset; nevertheless, it showed high support in both the coalescent (Figure 3A) and ML (Figure 2A) trees. The plastome with CDS regions (Figure 3B) and the CDS regions without paralogous genes (Figure 3C) recovered identical and nearly identical topologies to the plastome SNP trees with only biallelic loci and without indels (Figure 3D) and with 10% missing data (Figure 3E), which, in turn, recovered topologies very similar to those obtained from nuclear data with the same two filters (Figures 3I and J) and with a linkage disequilibrium (LD) filter (Figure 3K), but disagreed with the topologies obtained from mitochondrial data, both with high support. However, in the ML trees, many branches showed higher support (Figures 2G and H). Moreover, the mitochondrial trees based on CDS regions were discordant regardless of the inference method.

Using ML methods, the plastome trees (Figures 2A–E) were very similar, except for the without indels tree, which was identical to the plastome without indels coalescent tree and displayed high support for most branches. The mitogenome trees (Figures 2F–H) resembled the nuclear SNP tree, but some discordances were present (Figures 2I–K). A consistent separation between *S. jussiaei* and *S. aciculare* was observed in both coalescent and ML mitochondrial trees, whereas these species were grouped in the same clade (Clade E) in all other topologies, except for the plastome without indels (Figures 2A and 3A), which was identical to the plastome no-indel coalescent tree (Figure3A) and had high support in both. Among the mitochondrial trees (Figures 2F–H), the CDS-based topology (Figure 2F) was the most discordant and contained many low-support branches, while the mitochondrial SNP trees were identical. The three nuclear trees (Figures 2I–K) were identical, their clade groupings had high support, and they were similar to the CM trees.

Trees reconstructed from SNPs exhibited the least discordance, except for the mitochondrial coalescent trees, whereas mitochondrial trees reconstructed from CDS regions were highly discordant regardless of the inference method, not only in relation to other topologies but also among themselves. Furthermore, the presence or absence of indels appeared to have some effect on the topological differences among the other trees, although they remained largely similar to one another.

Although the bootstrap provides high support across all branches of the topologies inferred with the coalescent method, there is a variation in support among the ML trees, with some topologies displaying branches with low support (Figure 2F, H, J). In these cases, site concordance factors (sCFs) generally mirror the low support observed in certain branches, such as in the mitochondrial dataset restricted to CDS regions (Figure 2F). The clustering of clades receives substantial sCF support in most ML trees, with the exception of clade C, composed of *S. vaillantii* and *S. stramonifolium*, which shows low support in some topologies (Figure 2B, C). Nevertheless, all clades (A, B, C, D, and E) exhibit reduced sCF support in the mitochondrial trees (Figure 2F–H), and many internal branches in these same trees are also characterized by low support.

## DISCUSSION

### What could be recovered with shallow genome skimming?

In Straub et al. (2012), a coverage of approximately 30× using genome skimming was sufficient to recover high-quality rRNA sequences, and 30× was also adequate for assembling the complete plastome. However, the authors encountered difficulties in assembling the mitogenome, with several missing regions. This result is similar to ours, as we were unable to assemble the mitochondrial genome using genome skimming, although we did obtain recovery for multiple genes (Appendix S5).

Accurate assembly of the mitogenome in plants is hampered by the presence of highly enriched repetitive sequences and frequent recombination. Because Illumina read lengths often do not span longer repeats, these regions may not be completely assembled (Wu et al., 2020).

Although previous studies (e.g., Straub et al., 2012; Ripma et al., 2014) successfully retrieved low-copy nuclear genes for the COSII gene set, we did not obtain the same result consistently, though we recovered some copies (Appendix S2). A coverage of 10× was sufficient to recover high-quality single-copy nuclear regions in *Vitis* (Vitaceae) using deep genome skimming (Liu et al., 2021). However, even when filtering our data to a coverage range between 5× and 100×, we did not achieve the same success.

The predominance of the plastome as the primary target in genome skimming approaches results from a combination of technical and biological factors: i) the plastome occurs in a high number of copies per cell, often in the hundreds to thousands, leading to disproportionately high representation even when sequencing depth is relatively low (Straub et al.,, 2012; Nevill et al.,, 2020); and ii) the plastome has a highly conserved structure, and its structural stability facilitates the assembly of complete genomes from short reads, even from degraded samples, as shown in studies using herbarium material (e.g., Nevill et al., 2020).

Reasons for the lower use of the nuclear genome compared to the plastome in phylogenetic reconstructions may include shallow depth for the low-copy fraction of the genome, large genome sizes, and the trade-off between the two (Reginato, 2022), in addition to difficulties in complete recovery via genome skimming, as shown above, and structural complexity associated with frequent rearrangements and recombination (Moller et al., 2021).

It is possible that we failed to recover low-copy genes due to shallow coverage in these regions. Low-copy loci are already less represented than repetitive regions, and if coverage is low in these loci, they may fall below the threshold required for assembly. Despite these limitations, the SNP-based approach circumvented these difficulties by recovering relatively long matrices for phylogenetic inference (see Table 2).

### Topologies and putative causes of incongruence

The stability of groupings across most trees, especially Clades A and B, suggests that phylogenetic relationships within certain subgroups of spiny *Solanum* are robust and unlikely to change regardless of the genome used for inference.

Interestingly, several groupings observed in the plastome tree without indels (Figures 2A–, 3A–), under both inference methods, are uncommon when compared to the stable clade arrangements recovered in the other trees, yet these topologies are identical to each other. Paradoxically, these are precisely the groupings that receive the highest sCF support.

Although plastome trees without indels (Figures 2A, 3A) exhibit high bootstrap values and high sCF, some relationships are not congruent with the phylogenies currently available for the Leptostemonun clade (Stern et al., 2011; Gouvêa, 2022). For example, while *S. lycocarpum* and *S. leptostachys*, which belong to the same clade, are generally clustered together in most trees, in our analyses *S. lycocarpum* is instead grouped with *S. stramoniifolium*, a member of the Lasiocarpa clade. This could represent a cytonuclear discordance, but the dataset assembled here is not ideal for testing that.

Based on the studies of Levin et al (2006), Stern et al., (2011) and Gouvêa (2020), the relationships recovered in the phylogenies were largely expected. Clade A, for example, is composed of species belonging to the Crinitum clade recognized by Gouvêa, (2020). Clades A, B, C, D, and E were recovered in most phylogenetic reconstructions, although support values and relationships among them varied depending on the dataset and inference method employed. The relationship between clades A and B was particularly frequent in the maximum likelihood trees, being recovered in nine out of eleven analyses, whereas in the coalescent trees this relationship was recovered in only five out of eleven reconstructions.

Considering that the sCF metric relies exclusively on informative site patterns of maximum parsimony inferred from quartets, Kück et al. (2022) argue that both branch length and compositional heterogeneity can negatively influence its outcomes. Specifically, reference trees with long-branched taxa in close proximity or similar compositional states are likely to be overestimated, whereas more internal branches reflecting greater heterogeneity tend to be underestimated, thereby compromising the accuracy of the measure. In our study, the exclusion of indels may have reduced heterogeneity in the dataset, which likely resulted in the overestimation of external branches and the underestimation of internal branches, as observed in the plastome tree without indels (Figure 2A). This pattern is consistent with the limitations highlighted by the authors and suggests that the interpretation of high sCF support values, at least in this case, should be approached with caution.

We found cytonuclear discordance among topologies, a common event in plant phylogenomics (Morales-Briones et al., 2018; Qin et al., 2024). However, our results indicate that nuclear SNP and plastome SNP trees are similar, while cytonuclear discordance is more pronounced in the mitochondrial trees (Figures 2F–H; Figures 3F–H). Although the CDS-based tree inferred with coalescent methods received high support, the ML tree with the same regions received only low to moderate support. Dominicus et al. (2025) termed the discordance between mitogenome and plastome “mitoplastomic discordance” after also finding strong incongruence between these genomes. In our study, this discordance was greater in coalescent inference.

Dominicus et al. (2025) also argued that mitochondrial and plastid maternal inheritance might not be as strict as previously thought, and that lateral transfer may introduce foreign sequences, helping explain mitoplastomic discordance. Maternal inheritance is the most common pattern for mitochondria and plastids, but uniparental-paternal and biparental inheritance have also been documented in flowering plants (Rose, 2019; Sakamoto and Takami, 2024). For instance, low temperatures in *Nicotiana tabacum* increase paternal plastid inheritance (Chung et al., 2023), and biparental mitochondrial inheritance has been documented in *Pelargonium* (Weihe et al., 2009).

Although CDS regions were useful for producing consistent plastome phylogenies with both inference methods, mitochondrial phylogenies were highly incongruent with the plastome. Given that mitogenomes can exhibit heterogeneous evolution, recombination, and gene transfers, incongruences are to be expected. Nevertheless, studies such as Lin et al. (2025), analyzing 41 mitochondrial CDS in 481 angiosperm species across 335 families and 63 orders, obtained moderate to high bootstrap support (>70%), but reported limitations in groups such as rosids and asterids (the latter including Solanaceae), suggesting that the performance of these regions depends on the evolutionary context of the taxon.

We faced difficulties in recovering the complete mitogenome, so missing data in CDS regions could be causing the incongruence or phylogenetic imprecision in the mitochondrial trees. Even though we recovered these regions, they were not recovered uniformly. Another factor potentially contributing to mitochondrial tree incongruence is the occurrence of structural rearrangements, as mitogenomes are known for high plasticity and frequent recombination in flowering plants (Moller et al., 2021). These incongruences are more evident in mitochondrial ML trees.

It is also important to note that our aim was not to produce a complete phylogenomic inference for South American Leptostemonum lineages, but to test the utility of SNPs recovered from shallow genome skimming for phylogenetic inference. Our taxon sampling is therefore limited, and Nabhan and Sarkar (2011) demonstrated that sparse taxon sampling can negatively impact phylogenetic reconstruction. However, coalescent approaches are less affected by sparse taxon sampling compared to other inference methods (Bravo et al., 2019), which may explain why mitochondrial ML trees showed greater incongruence, while coalescent trees disagreed little with other datasets, in line with Bravo et al. (2019).

This incongruence pattern may be related to factors such as incomplete lineage sorting (ILS) and physical linkage among markers. The CDS set is smaller compared to the nuclear genome (or associated with fewer genes) and is physically linked, leading to unitary inheritance despite the possible influence of multiple copies on evolution. This increases the likelihood that the phylogenetic signal reflects the specific history of the organelle (including capture, hybridization, or drift) rather than the species history. In contrast, when using nuclear SNPs filtered to minimize LD, the risk of results being influenced by physical linkage or persistent ILS is reduced.

Furthermore, the impact of applied filters, especially for LD and missing data, should be considered with caution, as overly restrictive filters may reduce noise but also eliminate valuable phylogenetic signals (Stull et al., 2023; Suissa et al., 2024). This aligns with our observation that the mitochondrial ML CDS tree does not match the plastome CDS tree resolution.

Finding discordance in our study is not surprising. One of the most recent large-scale analyses of *Solanum* (Gagnon et al., 2022) found polytomies in the genus due to putative ILS. Gagnon et al. (2022) argued that ILS in *Solanum* is driven by rapid diversification in parts of the phylogeny. Echeverría-Londoño et al. (2020) found significant heterogeneity in diversification rates along the branches of a *Solanum* phylogeny, with a global mean of 0.25 lineages Myr⁻¹. The Leptostemonum clade was where the highest diversification rates estimated by Echeverría-Londoño et al. (2020) lie.

However, Messeder et al. (2024), using 1,786 low-copy nuclear genes for 247 *Solanum* species, showed that increasing the dataset resolved polytomies and uncertainties identified by Gagnon et al. (2022), suggesting that ILS or linkage is associated with organellar genomes, supporting our hypothesis above. Nevertheless, the work of Messesder et al. (2024) do not explore or discuss discordances found and seem only to have advanced in the idea that the more data the better. Although the approach of Messeder et al. (2024) of having full transcriptomes for phylogenetic inference might be seen as ideal, its implementation in larger scales might be limited by budget constraints.

Cytonuclear discordance between plastid and nuclear trees is also widely reported. Liu et al. (2025) used complete plastome data and nuclear SNPs to reconstruct the phylogeny of *Lithocarpus* (Fagaceae) and found significant incongruence between the phylogenies. ILS was cited as contributing to phylogenetic discordance, although later studies attributed cytonuclear discordance to introgression (Lee-Yaw et al., 2018). Although we did not test for introgression signals, historical introgression is plausible given the extensive record of the phenomenon in *Solanum* (e.g. Labate and Robertson, 2012; Hardigan et al., 2017).

Coverage uniformity may also play a role in incongruences. Jenke et al. (2022) found a correlation between low uniformity and the presence of ambiguous nucleotides. Eaton et al. (2017) showed that in *Viburnum* (Adoxaceae), insufficient or uneven coverage accounted for a portion of missing data. In their study, doubling the coverage nearly doubled the number of informative sites and increased the number of loci with shared data by over tenfold. Jenke et al. (2025) also showed that the sequencing platform influences coverage uniformity. The usage of SNP approach nevertheless can be less affected by coverage uniformity, since the expected coverage can be controlled.

A bias in SNP acquisition from genome skimming, especially regarding nuclear genome representation, could also be contributing to the observed patterns. Nevertheless, we recovered a substantial number of SNPs (Table 2). We obtained up to 138,560 nuclear SNPs (without data restrictions), a value close to the lower range reported in *Carya* (Juglandaceae; ≈180,000 SNPs; Literman et al., 2022) using the same shallow genome skimming approach, and slightly exceeding that reported in Tillandsioideae (116,478 nuclear SNPs; Loiseau et al., 2021), which also included 5,171 plastome SNPs and 1,069 mitochondrial SNPs.

In Melastomataceae (Reginato et al., 2026), the recovery of low-copy nuclear genes using genome skimming was highly successful, in contrast to our results. However, while the phylogenomic inference in Melastomataceae relied exclusively on low-copy nuclear markers, our approach also captured mitochondrial and plastid markers, which may have hindered the recovery of low-copy nuclear genes due to the predominance of organellar regions in the sequencing output, as previously reported in previous studies ( Liu et al., 2021).

Our findings further highlight the limitations of genome skimming as a standalone approach for phylogenomics, particularly in assembling complete mitogenomes and evenly capturing low-copy nuclear genes. For broader *Solanum* studies, it may be advisable to combine genome skimming with targeted hybridization capture techniques such as Hyb-Seq, which have shown greater efficiency in recovering low-copy/single-copy nuclear loci (Weitemier et al., 2014; Johnson et al., 2019; Straub et al., 2020; Xu et al., 2025). However, Liu et al. (2021), demonstrated that deep genome skimming in *Vitis* was as effective as Hyb-Seq for capturing large SCN datasets.

Despite the limitations of genome skimming, our results underscore the utility of SNPs derived from this approach for phylogenetic inference. The recovery of long, informative matrices allowed for phylogenetic reconstructions with high support across multiple clades, partially overcoming the difficulties associated with recovering complete nuclear genes and providing genome-wide universal markers.

Similar strategies have been successful in other taxa with analogous genomic complexity, such as *Lachemilla* (Rosaceae; Morales-Briones et al., 2018), *Viburnum* (Adoxaceae; Eaton et al., 2017), and Vitaceae (Liu et al., 2021), suggesting that SNP-based approaches can be a practical and efficient means to resolve phylogenies, particularly when genome skimming cannot produce homogeneous datasets.

Based on our findings, we recommend that future *Solanum* phylogenomic studies: i) ensure a minimum coverage of 30× for target regions, especially organellar ones; ii) adopt SNP-calling pipelines that carefully consider linkage disequilibrium and missing data; and iii) conduct a preliminary assessment of coverage uniformity before phylogenetic inferences, as outlined by Suissa et al. (2024).

## Conclusion

In summary, our study demonstrates that while genome skimming is effective for recovering organellar genomes and generating informative SNPs, it is limited in assembling mitogenomes in *Solanum*. Despite limitations in recovering nuclear regions, particularly low-copy loci, the SNP matrices derived from genome skimming proved highly informative, producing stable, congruent, and well-supported phylogenies. Our findings indicate that, even with variable coverage and uniformity, genome skimming can be a powerful tool for recovering SNPs and using them in strongly supported phylogenetic inferences. However, to obtain a more complete and homogeneous dataset, particularly for low-copy nuclear loci, deep genome skimming or a combination of genome skimming with targeted capture methods such as Hyb-Seq, along with careful SNP filtering and prior coverage evaluation, may be more appropriate.

## Author Contributions

LLG, JD, and PP conceptualized the manuscript; YFG and LLG collected samples and generated data; PP, JD and LLG performed analysis; RTLA and JD draw the final figures; writing was led by RTLA and LLG with contributions from all authors.

## Acknowledgements

We thank Clarisse Palma Silva, Ingridy Moura, Thiago J.C. André, and Thaís E. Almeida for the support with labwork. Juan D. Tovar contributed samples and shared useful R scripts. The authors thank CNPq (grants 422191/2021-3, 408914/2023-8, and 404996/2024-8 awarded to LLG and 152961/2024-0 awarded to YFG), RTLA acknowledges FAPESQ/PB and SECTIES for the scholarship granted through Call No. 11/2024 – Granting of Sandwich Master’s and Sandwich Doctorate Scholarships for International Mobility under the “Paraíba sem Fronteiras” Program, which resulted in this text as one of the products from the period spent at the University of Helsinki, Finland. The author also acknowledges the regular scholarship granted by FAPESQ-PB through Call No. 08/2023. The CSC – IT Center for Science, Finland, provided free access to high-performance computing resources essential for conducting this research in support of academic study and education. We also thank the support of the TFK - MOMENT project and the support of Helsinki University Library for open access publication.

## Supporting Information

Additional Supporting Information may be found online in the Supporting Information section at the end of the article

S1-Attempt to recover genes using the A353 bait set in HybPiper

S2- Attempt to recover genes using COSI.

S3- Attempt to recover genes using COSII.

S4- Heatmap showing percentage length recovery of the rRNA gene across samples

S5- Heatmap of the tentative capture of mitochondrial genes

